# Spinach (*Spinacia oleracea* L.) Flavonoids Are Hydrolyzed During Digestion and Their Bioaccessibility Is Under Stronger Genetic Control than Raw Material Content

**DOI:** 10.1101/2025.09.27.678982

**Authors:** Michael P. Dzakovich, Alvin L. Tak, Elaine A. Le, Rachel P. Dang, Benjamin W. Redan, Geoffrey A. Dubrow

## Abstract

Spinach (*Spinacia oleracea* L.) is a commonly consumed crop with a diverse array of unique flavonoids. These molecules likely contribute to the health benefits associated with spinach consumption. However, little is known about the genetic diversity of these molecules, their bioaccessibility, and the heritability of these traits. We assembled a diversity panel of 30 F_1_ and open pollinated spinach accessions and cultivated them under controlled conditions over two periods. Quantification of 39 flavonoids revealed that their concentration is largely influenced by environmental factors and at least two divergent branches in the flavonoid biosynthesis pathway may exist. Despite generally similar trends in the amounts of major flavonoids, open pollinated and F_1_ varieties of spinach could be distinguished based on the concentrations of minor flavonoid species. Broad sense heritability estimates for absolute bioaccessibility accounted for more genetic variation than raw material content, suggesting that this trait is preferable for breeders seeking to alter the phytochemical profile of spinach. Lastly, we found that several spinach flavonoids are unstable under digestive conditions, made evident by the proportion of aglycones rising from 0.1% to approximately 15% of total flavonoids after digestion. Together, these data suggest that spinach flavonoid biosynthesis and bioaccessibility are complex and contextualizes how these molecules may behave *in vivo*.

## 1. Introduction

Flavonoids are among an array of phytochemicals associated with positive health benefits from fruit and vegetable-rich diets. They have been hypothesized to have antioxidative, anti-inflammatory, and anticarcinogenic properties *in vivo* [1–3]. Spinach (*Spinacia oleracea* L.) is a dietary source of flavonoids with significant concentrations reported up to and beyond 224 mg/100 g fresh weight [4–7]. While concentrations are high, the profile of flavonoids in spinach departs from what is typically found in fruits and vegetables. Spinach flavonoids feature a patuletin, spinacetin, or jaceidin backbone that is further modified by glycosylation, glucuronidation, and conjugation with phenolic acids [6,8]. Although little is known about the specific roles of spinach flavonoids on human health, they remain a trait of interest due to their structural similarity to more common dietary flavonoids [9,10].

Plant breeding programs tend to focus on increasing nutrients and/or phytochemicals in crops. However, increased amounts of a given substance in a plant does not guarantee greater bioefficacy. Food matrix factors such as protein, starch, and fiber can affect the release of a compound from the food (*bioaccessibility*) and the absorption into the body (*bioavailability*) [11,12]. Bioaccessibility and bioavailability are becoming key considerations in breeding programs focusing on mineral elements in rice [13], wheat [14], common bean [15], maize [16], and sorghum [17]; carotenoids in maize [18], banana [19], and spinach [20]; as well as polyphenols in blueberry [21]. While randomized clinical trials in humans are considered the gold standard to assess the impact of either foods or molecules on health outcomes, they are costly and logistically challenging to operate. Bioaccessibility estimates from *in vitro* models of digestion associate well with human clinical data and are frequently used as a cost-effective surrogate to estimate bioavailability [22,23]. These models are crucial for efficiently phenotyping populations of plants for plant breeders to make selections that might ultimately be used in clinical applications.

Only limited reports in the context of post-harvest processing exist for spinach flavonoid bioaccessibility. Freezing spinach preserves flavonoid profiles while modestly enhancing bioaccessibility; ostensibly through changes in the food matrix [24]. Processing techniques such as spray drying and mixing with a protein carrier have also been shown to modulate spinach flavonoid bioaccessibility in a processing-dependent manner [25]. There are currently no data on the role of genetic background on flavonoid bioaccessibility in spinach and if this trait is heritable in this crop. We hypothesized that spinach flavonoid bioaccessibility is heritable and a distinct trait from raw material flavonoid content.

The primary goals of this study were to define the range and concentration of flavonoids in a diverse population of spinach using a comprehensive ultra-high performance liquid chromatography mass spectrometry (UHPLC-MS/MS) method, determine the bioaccessibility of spinach flavonoids using a three-stage static *in vitro* digestion model coupled with UHPLC-MS/MS, and to determine the broad sense heritability of spinach bioaccessibility. These data would serve as a resource to plant breeders seeking to optimize the potential health benefits of spinach and to allow for the development of unique germplasm that can be tested in clinical settings.

## 2. Materials and Methods

### 2.1. Chemicals and reagents

Reagents purchased from Sigma Aldrich (Sigma Chemical Co., St. Louis, MO, US) included quercetin-3-glucoside (99%), naringin (99%), naringenin (99%), mucin, alpha-amylase (lot: SLCM1439), pepsin (lot: SLCG6556), lipase (lot: SLCJ9303), pancreatin (lot: SLCK9806), and bile (lot: SLCJ7934). Urea, uric acid, potassium chloride, sodium sulfate, sodium phosphate, sodium chloride, sodium bicarbonate, LC-MS grade water, acetonitrile, methanol, and formic acid were purchased from Fisher Scientific (Fisher Scientific, Waltham, MA, USA). Patuletin (99%) was purchased from Extrasynthese (Extrasynthese, Genay, France) and jaceidin (99%) and spinacetin (99%) were purchased from Key Organics (Key Organics, Camelford, United Kingdom). Taxifolin (99.99%) was purchased from Selleck Chemicals (Selleck Chemical LLC, Houston, TX). StrataX polymeric reversed phase solid phase extraction (SPE) cartridges (30 mg, 33 µm pore size, 1 mL volume) were purchased from Phenomenex (Torrance, CA).

### 2.2. Germplasm Selection and Growth Conditions

Thirty spinach accessions were selected to maximize genetic variation in our population based on previous sequencing and phenotyping efforts [20,26]. Accessions previously genotyped by Qin and colleagues were selected from unique sub-populations separated by sequence diversity as determined by STRUCTURE2 and MEGA6 [27,28]. Germplasm previously characterized by Hayes and colleagues was selected for this study based on variation in carotenoid bioaccessibility. Additional metadata for each accession can be found in Supplemental Table 1.

Spinach plants were cultivated according to Qin and others with modifications described by Dzakovich and colleagues [6,26]. For each cultivar, seedlings were thinned to six individual plants per one gallon pot. Pots were randomly distributed throughout a PGW36 walk-in growth chamber (Conviron; Winnipeg, Canada) maintained at 300 μmols/m^2^/s photosynthetically active radiation for 12 hours per day, 50□±□10% relative humidity, and 20□±□0.5°C/15□±□0.5°C day and night temperatures, respectively. Plant positions were re-randomized each day to mitigate positional differences in environmental factors such as light intensity. Plants were considered mature at the six to eight leaf stage (five to six weeks after sowing), edible tissues harvested, and stored at -80 □ until analysis. This population was grown and harvested as described above two times.

### 2.3. Sample processing

Frozen tissue was homogenized 1:1 with MilliQ water for approximately one minute using a VWR 250 Homogenizer, 10032-766 (Radnor, PA, USA). Homogenate was immediately aliquoted for raw material analysis, *in vitro* digestion, and dry matter content. Homogenates were stored at -80 □ until use. All subsequent analyses were conducted in triplicate.

### 2.4. In vitro digestion

Spinach samples were subject to a three-phase static *in vitro* digestion method described previously for food products including spinach and sorghum [20,29,30]. In oral phase, spinach homogenate (0.4 g) was mixed with 1.2 mL of simulated saliva containing 31.8 mg/mL alpha-amylase, incubated at 37 □ for 10 min at 120 oscillations per minute (OPM). During the gastric phase, 0.4 mL of 10 mg/mL pepsin solution in 0.1 M HCl was added and samples were further diluted with 2.7 mL of saline (0.9% NaCl). Sample pH was adjusted to 2.5 using 1.0 M HCl. Saline was added to dilute samples to 8 mL and sample headspace was replaced with nitrogen gas prior to incubation at 37 □ for 1 hour at 120 OPM. Sample pH was then adjusted to 5.0 using 1.0 M NaHCO_3_ to initiate the intestinal phase. Pancreatin-lipase solution (0.4 mL; 20 mg/mL) in 0.1 M NaHCO_3_ and porcine bile extract (0.6 mL; 30 mg/mL) were added and sample pH was adjusted to 7.0. Sample volumes were adjusted to 10 mL using saline prior to an additional blanketing with nitrogen gas. Samples were then incubated at 37 □ for 2 hours at 120 OPM. At the end of intestinal phase, digesta was aliquoted and 4 mL were stored at -80 □ for later analysis. The remaining digesta were blanketed with nitrogen gas and centrifuged at 3428 x *g* for 75 minutes at 4 □ (Eppendorf 5920R). Approximately 4 mL of each aqueous fraction was filtered through 0.22 µm cellulose acetate filters and reserved to quantify bioaccessible flavonoids.

### 2.5. Extraction of flavonoids from fresh spinach and aqueous fractions

Flavonoids were extracted in duplicate as previously described by Dzakovich and others [6]. Dried spinach flavonoid extracts were redissolved in 5 mL of 1:1 methanol:water + 0.1% formic acid containing 4% 20 µM taxifolin as an internal standard to correct for variation in mass spectrometer response and filtered through 0.2 µm PTFE filters prior to analysis. Flavonoids from aqueous fractions were semi-purified using solid phase extraction. Aqueous fractions (500 μL) were pipetted into 3 mL Strata-X 33 µm, 30 mg polymeric reversed phase SPE cartridges (Phenomenex) and diluted with 2500 μL of 0.1% formic acid in water. Samples were loaded under vacuum and rinsed with one volume of 0.1% formic acid in water followed by one volume of 1.0% formic acid in water. Cartridges were dried under vacuum for approximately 10 minutes and rinsed with one volume of hexanes to remove nonpolar and semi-polar impurities (e.g., carotenoids). Cartridges were dried for 2 minutes under vacuum and analytes were eluted with 2 mL of methanol with 0.1% formic acid. Vials of eluent were dried under a stream of nitrogen gas and resolubilized in 500 μL of 1:1 methanol:water with 0.1% formic acid and 4% 20 µM taxifolin as an internal standard. Samples were passed through 0.2 µm PTFE filters and into LC vials prior to analysis.

### 2.6. Recovery estimates from SPE

Separate spike addition experiments were conducted to optimize the recovery of flavonoids from aqueous fractions. Spinach-free digesta was created using the *in vitro* digestion protocol detailed above and spiked with jaceidin, patuletin, spinacetin, naringenin, and naringin. Various washing and elution conditions were tested using the methods of Mengist and colleagues as a starting point [21]. Methanol with 0.1% formic acid produced the most consistent and high yielding results with estimated recoveries of 87%, 91%, 111%, 98%, and 99% for jaceidin, patuletin, spinacetin, naringenin, and naringin, respectively.

### 2.7. Analysis of spinach flavonoids

Flavonoids from both spinach fresh tissue and aqueous extracts were analyzed using an UHPLC-MS/MS method developed by Dzakovich and colleagues [6]. Briefly, flavonoids were separated on a Waters Acquity BEH C18 2.1 x 100 mm (1.7 µm particle size) column using a gradient of 0.1% formic acid in water (A) and 0.1% formic acid in acetonitrile (B). Column eluent flowed into a TSQ Altis tandem mass spectrometer (ThermoFisher) and spinach flavonoids were quantified against the external standard quercetin-3-O-glucoside when an authentic standard was not available. Optimized multiple reaction monitoring (MRM) experiments for 39 spinach flavonoids, including isomers, maximized signal to noise of each analyte. Dwell times were automatically adjusted to maintain 12-15 points per peak and analyte signal was corrected for using taxifolin. Spinacetin was excluded from analyses due to being below the limit of quantification for all samples and sample types.

### 2.8. Statistical Analysis

Plants were grown in a randomized complete block design with two replications in time and analyzed with the following model parameters:

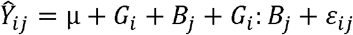

Where *Ŷ*_*ij*_ represents an analyte estimate; μ is the population mean for an analyte; *G*_*i*_ represents genetic factors; *B*_*j*_ represents the contribution from block; *G*_*i*_:*B*_*J*_ represents the interaction between genotype and block; and ℰ_ij_ is the residual error.

All statistical analyses and data visualization were performed using R statistical software [31]. Linear modeling including the calculation of best linear unbiased estimates (BLUEs) and random effects modeling was conducted using the package ‘lme4’ [32]. The restricted maximum likelihood (REML) was used to estimate variance components to calculate broad sense heritability [32]. Correlations were calculated and visualized using the ‘Corrplot’ package [33] while principal components were calculated using ‘FactoMineR’ and visualized with ‘ggplot2’ [34,35]. Hierarchical clustering to determine chemical similarity using Tanimoto coefficients was conducted with the package ‘ChemmineR’ [36].

## 3. Results and Discussion

### 3.1. Spinach flavonoid profiles are diverse, but dominated by five molecules

To determine the range and diversity of spinach flavonoid raw material content and bioaccessibility, we grew 30 unique accessions comprised of both F_1_ varieties and open-pollinated accessions over two periods of time. Approximately 56.5% of spinach flavonoids in raw material were patuletin derived while the remaining 31.6%, 6.7%, and 5.3% were derived from methylenedioxyflavones, spinacetin, and jaceidin. Concentrations in raw material ranged from 64.3 to 453.7 mg/100 g fresh weight (FW) with an average concentration of 185.8 mg/100 g FW (Supplemental Table 2). On average, approximately 67% of the total flavonoid profile was represented by five analytes including 5,3’,4’ -Trihydroxy-3-methoxy-6:7-methylendioxyflavone-4 - β-D-glucuronide (519.08 *m/z*), Spinatoside (521.09 *m/z*), Patuletin-3-O-β-D-2-ρ-coumaroyl glucopyranosyl-(1 → 6)-β-D-glucopyranoside (801.21 *m/z*), 5,4 -Dihydroxy-3,3 - dimethoxy-6:7-methylendioxyflavone-4 - β-D-glucuronide (533.09 *m/z*), and Patuletin-3-O-β-D-(2-feruloyl glucopyranosyl)-(1 → 6)-[β-D-apiofuranosyl-(1 → 2)]-β-D-glucopyranoside (963.24 *m/z*) (Supplemental Table 2). The profile and concentration ranges of spinach flavonoids in our population was consistent with previous reports [6,7,37,38]. Ranges of aggregated flavonoid classes presented on a log_10_ transformed y-axis can be found in Figure 1.

**Figure 1.**
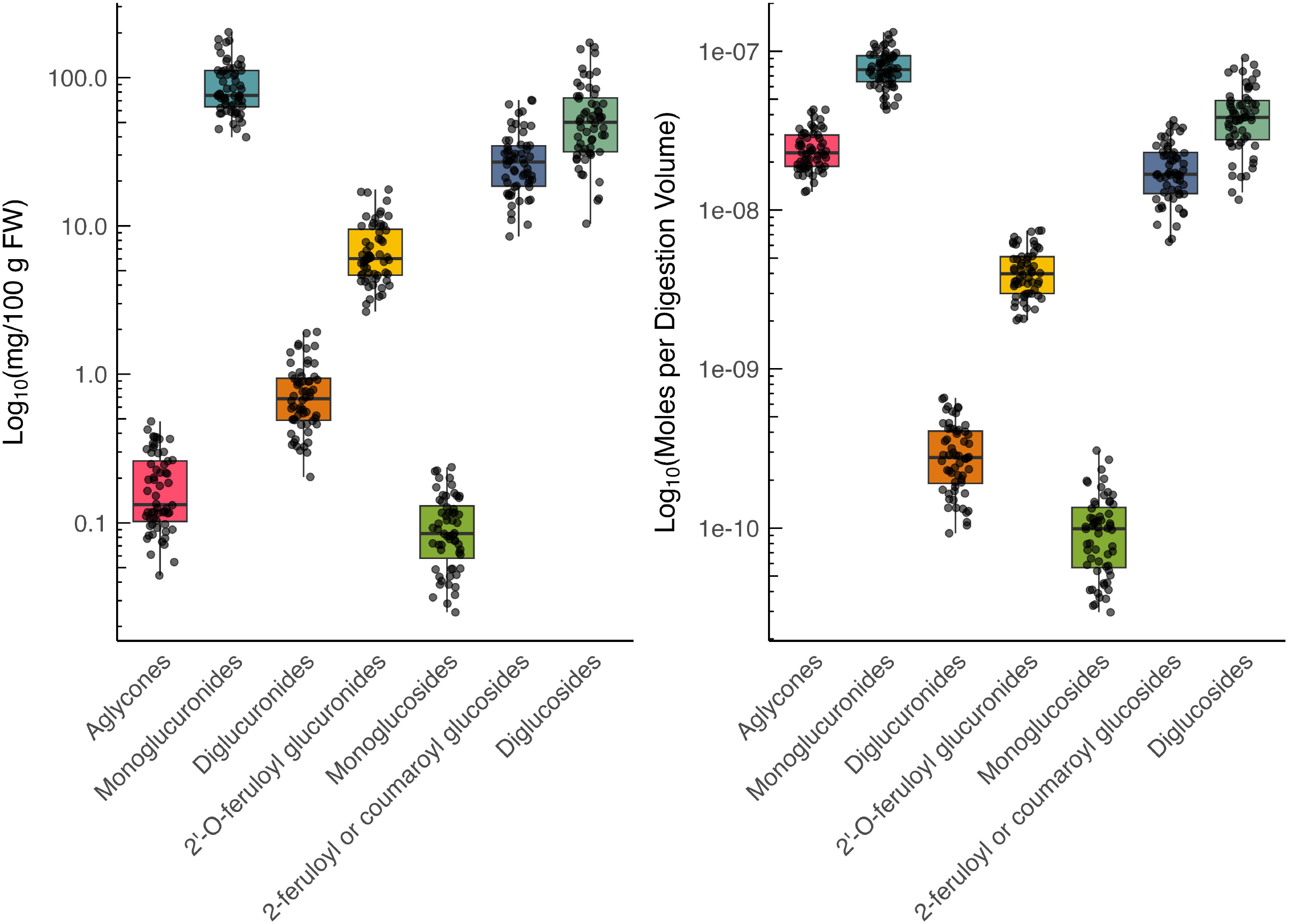
Box and whisker plots of aggregated spinach flavonoids in raw material (a) and aqueous fractions (b). Y-axes are log10 transformed to accommodate the wide range in concentrations calculated for these flavonoids. Each dot within a box and whisker plot represents an individual variety grown in block. Of note is the increase in relative abundance of aglycones after digestion (b).

### 3.2. Minor flavonoids drive differences between F_1_ and open pollinated spinach lines

Multivariate analysis using PCA revealed a distinction between open pollinated and F_1_ spinach varieties (Figure 2). Although principal component 1 explained a large proportion of total variance (53.4%, Figure 2), F_1_ varieties and open pollinated lines separated primarily on principal component 2 (17.95% total variation). Individual loadings for the input variables indicated that several minor flavonoids such as 5,4’-Dihydroxy-3,3’-dimethoxy-6:7-methylen-dioxyflavone-4’ -β-D-2′-O-feurloyl-glucuronide (709.14 *m/z*) likely contributed to the differentiation between F_1_ and open pollinated lines (Figure 2B). Although low in concentration relative to other flavonoids (0.53 – 6.62 mg/100 g FW), 709.14 *m/z* was on average 53.7% higher in F_1_ spinach varieties compared to open pollinated lines with similar trends for other variables (*p* < 0.001; Supplemental Table 3). Similar trends were true for other variables in the upper right quadrant of the loadings plot. These flavonoids may have been inadvertently selected for commercial germplasm. Flavonoids and other polyphenols have been reported to be influenced by human selection in other crops such as lettuce [39] and apple [40]. Alternatively, differences in minor flavonoids may be due to an overall increase in all flavonoids. Although statistically nonsignificant in our population (*p* > 0.05), our data shows that F_1_ spinach varieties were on average lower in total flavonoids compared to open pollinated lines (176.65 vs 192.93 mg/100 g FW; Supplemental Tables 2 and 3). Increases in minor flavonoids seen in F_1_ spinach varieties may be a consequence of lower competition for precursors and/or differences in enzymatic gene expression or enzyme efficiency. This phenomenon has been observed for carotenoids in crops such as maize [18,41], soybean [20,42], tomato [43] as well as polyphenols in apple [44], maize [45], and rice [46–48]. Additional follow-up experiments coupling next generation sequencing with an expanded population of wild and cultivated spinach are necessary to define the mechanism of this phenomenon in spinach.

**Figure 2.**
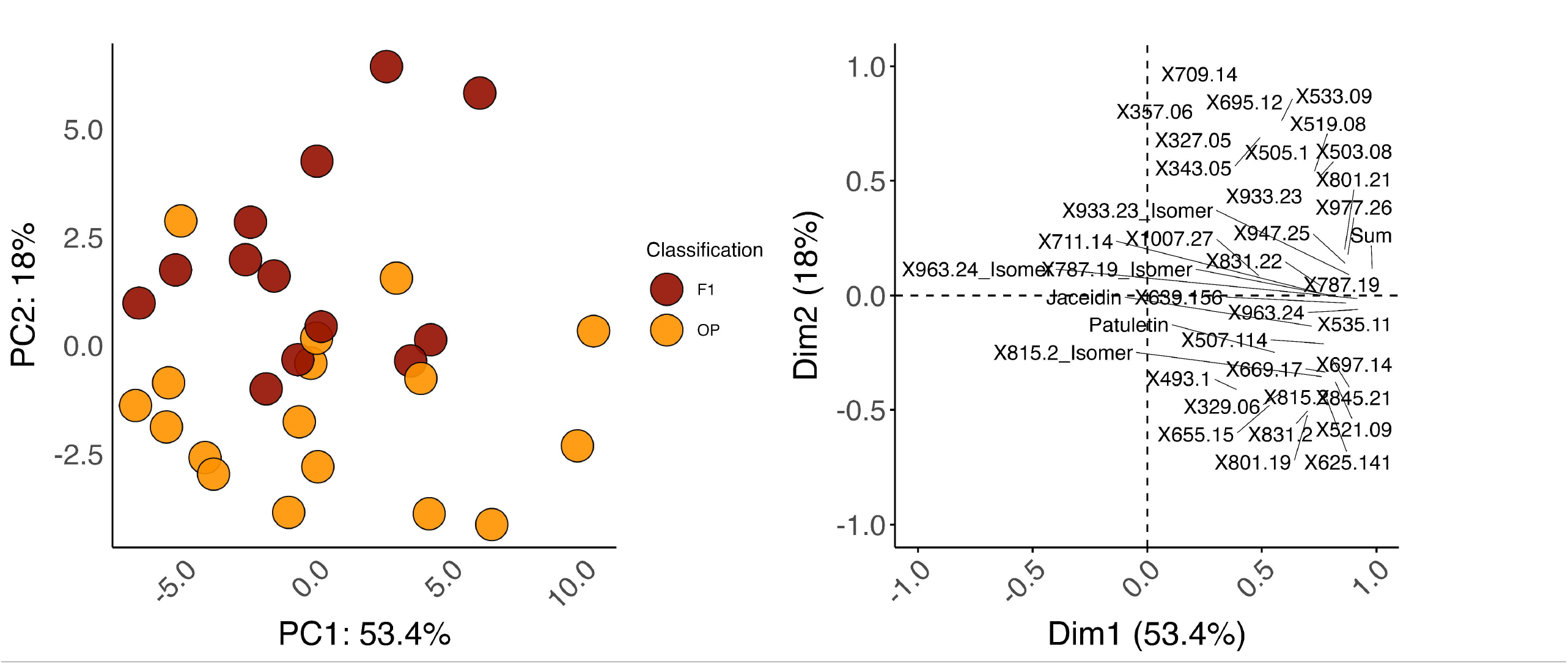
Scores (a) and loadings (b) plots from principal components analysis of raw material content. BLUEs for each analyte within a cultivar were utilized as input data to more accurately reflect genetic effects and minimize environmental distortion. “F1” indicates F_1_ varieties and “OP” indicates open pollinated lines.

### 3.3. Raw Material Content and Absolute Bioaccessibility Are Generally Associated, but Several Negative Associations Exist

To better understand the relationships between spinach flavonoids as well as raw content and absolute bioaccessibility, we performed correlation analyses (Figure 3). Matrices of Pearson’s correlation coefficients and *p*-values can be found in Supplemental Table 4. Generally, individual flavonoids were positively associated with each other indicating that concentrations rise and fell in concert. However, several analytes were negatively correlated with most other spinach flavonoids including 5,4’-Dihydroxy-3,3’-dimethoxy-6,7-methylenedioxyflavone (357.06 *m/z*), 5,4’-Dihydrox-3-methoxy-6,7-methylenedioxyflavone (327.05 *m/z*), 709.14 *m/z*, 695.12 *m/z*, 5,3,4 -Trihydroxy-3-methoxy-6:7-methylendioxyflavone (343.05 *m/z*), and to a limited degree 5,4 -Dihydroxy-3,3 -dimethoxy-6:7-methylendioxyflavone-4 - β-D-glucuronide (533.09 *m/z*) Figure 3. Interestingly, these are the same analytes that appeared to drive differences between F_1_ and open pollinated germplasm (Figure 2). Absolute bioaccessibility correlated well with raw material content (Figure 3) consistent with other reports in the literature [21,25,49–51]. A few analytes such as Spinacetin-3-O-β-D-glucopyranosyl-(1 → 6)- [β-D-apiofuranosyl-(1 → 2)]-β-D-glucopyranoside (801.19 *m/z*) displayed negative associations with the analytes mentioned above (e.g. 709.14 *m/z*) both in raw content and absolute bioaccessibility. Raw data for relative and absolute bioaccessibility can be found in Supplemental Tables 5 and 6, respectively.

**Figure 3.**
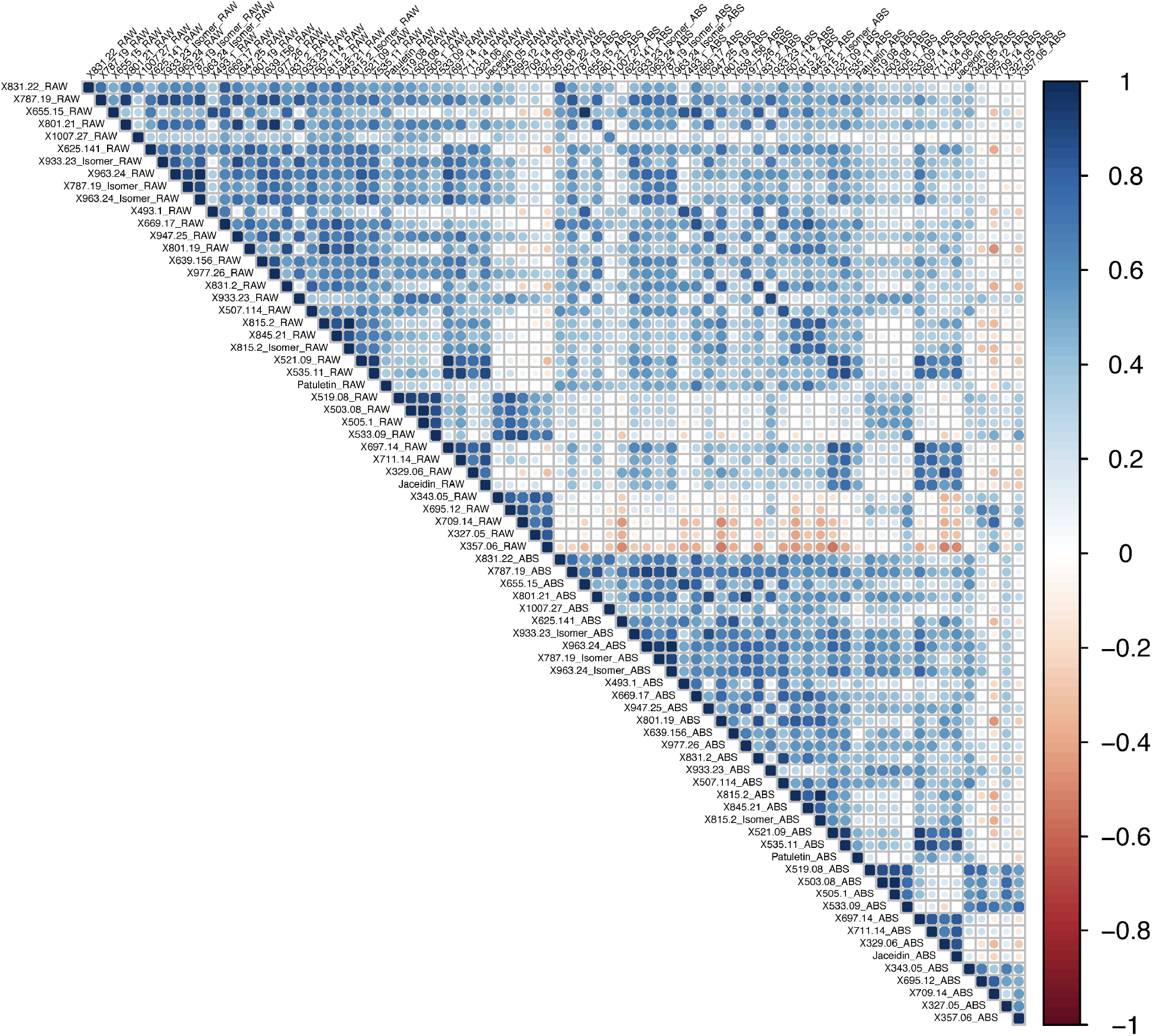
Correlation plot of raw material content and absolute bioaccessibility of the analytes measured in this study. Dark blue to dark red indicates a Pearson correlation coefficient spanning from 1 to -1, respectively.

Given that the analytes with negative correlation coefficients are methylenedioxyflavones, we hypothesize that the trends we observed are due to a split in the spinach flavonoid biosynthesis pathway whereby select methylenedioxyflavones compete for precursor molecules that feed into multiple segments of the pathway. While there have been strides to characterize the flavonoid profiles of spinach [6–8], the structural and regulatory aspects of this pathway remain largely unknown. Using Tanimoto coefficients, we sorted analytes profiled by our method using hierarchical clustering (Figure 4). Structures and SMILES abbreviations of each analyte in our UHPLC-MS/MS method can be found in Supplemental Table 7. These data were used in determining groups of aggregated flavonoids presented here (Figure 1; Supplemental Table 2). Clustering organized analytes by structural motifs and may suggest biosynthetic relationships among different spinach flavonoids. Similarity based on structural motifs may indicate common enzymes that act on multiple substrates. Additional studies leveraging genomic tools and techniques such as virus-induced gene silencing are necessary to definitively map the spinach flavonoid biosynthesis pathway.

**Figure 4.**
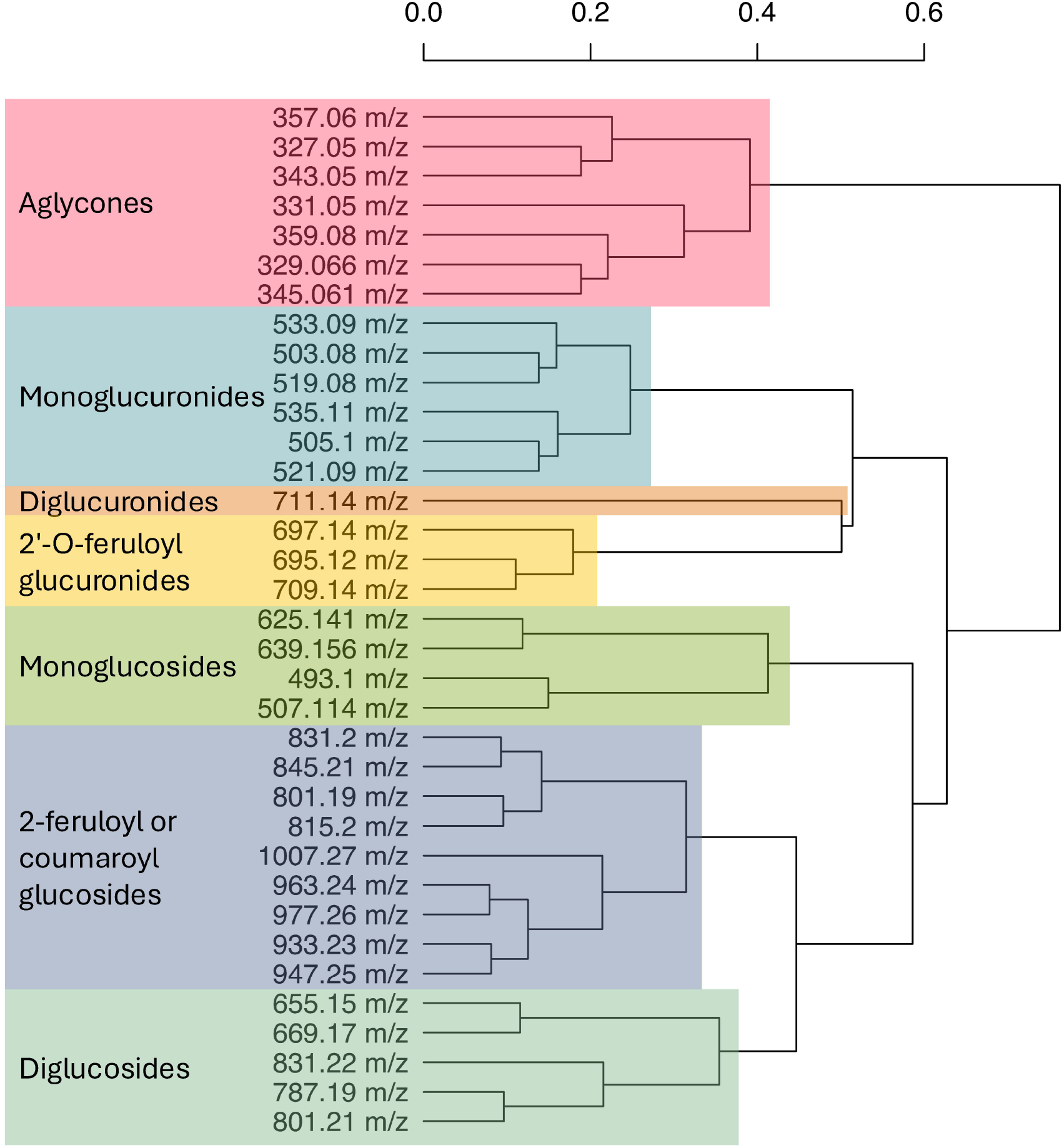
Dendrogram grouping spinach flavonoids based on Tanimoto coefficients. The scale bar above indicates Euclidean distance.

### 3.4. Spinach Flavonoid Absolute Bioaccessibility is More Heritable than Raw Material Content

To define the contribution of genetics relative to environmental factors, we calculated broad sense heritability on a line-mean basis for raw material content, relative bioaccessibility, and absolute bioaccessibility [52]. A graphical illustration of variance partitioning for model components cultivar, block, block:cultivar interaction, and residual can be seen in Figure 5. Variance for each model component for every analyte can be found in Supplemental Table 8. Broad sense heritability for raw material content ranged from 0 – 63% with an average of 22% (Supplemental Table 8). Contrary to our expectations, relative bioaccessibility exhibited extremely low estimates of broad sense heritability 0 – 31% with an average of only 2%. For virtually every analyte, most of the variation partitioned into either block or block:cultivar interaction. Reports of broad sense heritability estimates for flavonoids in spinach, or other leafy greens, are scant. A similar trend of moderately high estimates of broad sense heritability for relative content and absolute bioaccessibility, but extremely low for relative bioaccessibility, has been reported in blueberry [21]. Broad sense heritability estimates for quercetin-3-arabidonside, quercetin-3-glucoside, and total flavonoids in blueberry were all 10% or less for relative bioaccessibility, but substantially higher for raw material and absolute bioaccessibility [21]. These findings echo our own and highlight a disconnect between raw material content and absolute bioaccessibility.

**Figure 5.**
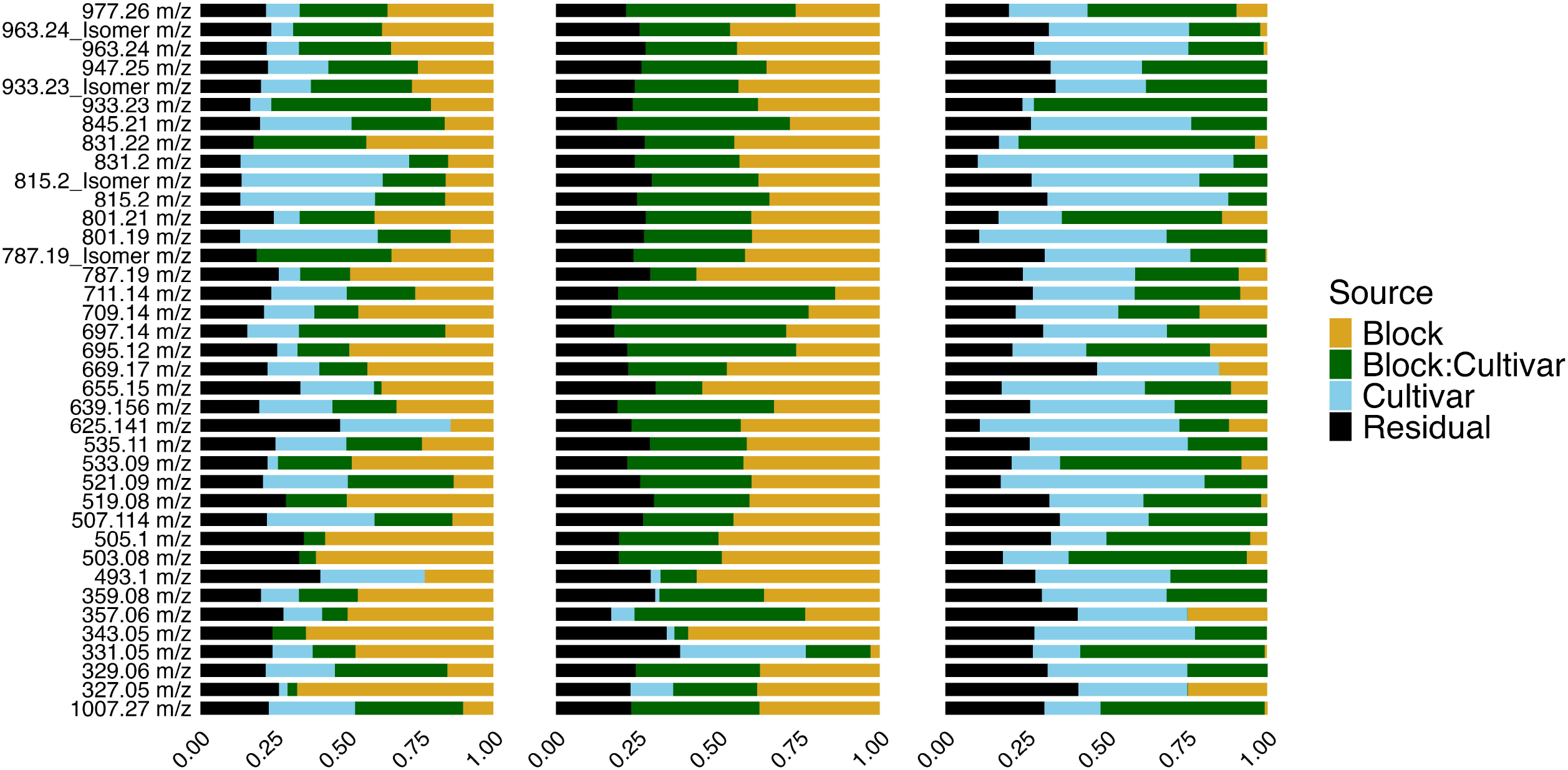
Variance partitioning estimates visualized for each flavonoid for raw material content (a), relative bioaccessibility (b), and absolute bioaccessibility (c).

Interestingly, absolute bioaccessibility, which is a product of raw material content multiplied b relative bioaccessibility, showed the highest estimates of broad sense heritability (Supplemental Table 8). Broad sense heritability for absolute bioaccessibility ranged from 4% – 76% with an average of 34%. Four analytes were estimated to be greater than 50% which included Spinacetin-3-O-β-D-(2-feruloyl glucopyranosyl) -(1→6)-[β-glucopyranosyl (1 → 2)]-β-D-glucopyranoside (625.141 *m/z*), (801.19 *m/z*), Patuletin-3-O-β-D-2-feruloyl glucopyranosyl-(1 → 6)-β-D-glucopyranoside (831.2 *m/z*), and 521.09 *m/z* (Supplemental Table 8). Only six analytes had broad sense heritability estimates that were equal to or less than raw material content, supporting the assertion that absolute bioaccessibility is under stronger genetic control than raw material content. This outcome is intriguing given that relative bioaccessibility was estimated near 0% for 30 out of 38 analytes measured.

Absolute bioaccessibility is a product of raw material content and relative bioaccessibility [53]. Our findings that absolute bioaccessibility heritability exceeded raw material heritability suggest that genetic control over traits related to digestion efficiency is stronger and more stable over flavonoid concentration. Similar findings have been reported for iron in maize [42] and carotenoids in spinach [20]. Heritability of absolute bioaccessibility may appear higher because absolute bioaccessibility combines the genetic differences in relative content with environmental variation in relative bioaccessibility, creating larger total differences in absolute bioaccessibility between genotypes. Even with low relative bioaccessibility, as is the case in our dataset, plants with slightly higher relative content tend to have higher absolute bioaccessibility on average, making absolute bioaccessibility more genetically predictable than relative content alone. Regardless, environmental factors tended to be a major influence in the phenotypes we measured.

While our data suggest that breeding for flavonoid content and profiles in spinach would be possible, but challenging, significant opportunities exist in controlled environment agriculture (CEA) to directly manipulate these traits by controlling plant growth conditions. Flavonoid biosynthesis is strongly controlled by light quantity and quality [54,55], temperature [56,57], and abiotic stress such as salinity [58]. Advances such as light emitting diodes (LEDs) allow for precise control of the light environment and other environmental parameters can be manipulated in modern CEA systems [59–61]. Additional research is necessary to determine how environmental conditions can be leveraged to fine tune the flavonoid profiles of spinach.

Although absolute bioaccessibility exhibited the highest broad sense heritability of nearly all phenotypes measured in this study, estimates tended to be relatively low compared to other crops [21,62– 64]. One possibility is due to the nature of spinach itself, which is a highly heterozygous, wind-pollinated outcrossing species [65–67]. Of the 30 accessions profiled in our study, 17 of which were open pollinated (Supplemental Table 1). Open pollinated accessions can breed true when cultivated under controlled conditions, but there is no guarantee of genetic uniformity seen in cultivated F_1_ varieties. As such, it is distinctly possible that some of our open pollinated, or heirloom accessions, many have been segregating for any number of traits. Our population structure may have had unaccounted for genetic variation that manifested as environmental variation. Future experiments can be conducted with larger populations comprised of cultivated F_1_ varieties to better understand the consequence of open pollination practices on flavonoid heritability. Regardless, heritability estimates for absolute bioaccessibility outperformed raw material content and this trait may be preferable for plant breeders to select against when considering potential health implications of their spinach germplasm.

### 3.5. Some Spinach Flavonoids Are Hydrolyzed During Digestion

Our estimates of relative bioaccessibility align with the limited reports in the literature that only monitored parts of the spinach flavonoid biosynthetic pathway [24,25]. Estimates of relative bioaccessibility theoretically range from 0 – 100%. However, several analytes in our study exceeded 100% such as 343.05 *m/z*, with relative bioaccessibility estimates ranging from 569.5 – 9489.1% (Supplemental Table 5). Over 99.9% of spinach flavonoids are O-glycosylated in raw spinach [6], yet this proportion shrinks to approximately 85% after digestion with a proportional rise in aglycones (Figure 1; Supplemental Table 2). Compared to C-linked glycosylated flavonoids, O-linked glycosylated flavonoids are considerably weaker and more prone to chemical hydrolysis [2]. *In vitro* digestions such as the one conducted in this study feature a 1-hour gastric phase at pH 2.0 at 37 □. While there is debate as to whether gastric conditions are sufficient to induce chemical hydrolysis [68,69], there is evidence to suggest that hydrolysis may occur with some polyphenols [21,70]. For example, Mengist and colleagues reported up to 4900% relative bioaccessibility for caffeic acid and over 100% relative bioaccessibility for catechin, epicatechin, flavon-3-ols, and total flavonoids. Given that some of these molecules have been found to be condensed with other analytes like anthocyanins [45], it is feasible that analytes reported at >100% relative bioaccessibility may be degradation products formed during digestion from more complex phytochemicals. Grace and colleagues did not report relative bioaccessibility over 100% for any of the 18 spinach flavonoids they tracked, but did report 533.09 *m/z* above 90% in some of their treatments [25]. In animals and humans, we would anticipate this conversion to occur at an even greater rate due to enzymatic activity at the brush boarder [71,72] and interactions with the gut microbiome [1,73].

*In vitro* digestions similar to the one conducted in this study provide a cost-effective way to screen germplasm for bioaccessibility, a measure that generally associates well with bioavailability for many molecules [11]. Analytes we observed to increase after digestion did so at the expense of other analytes. For example, 695.12 *m/z* had an average relative bioaccessibility of 22.6%, the structurally related 519.08 *m/z* at 20.6%, and the aglycone 343.05 *m/z* at 2953.6% (Supplemental Table 5). We hypothesize that the elevated appearance of 343.05 *m/z* was due to the successive loss of ferulic acid from 695.12 *m/z* and loss of glucuronic acid from 519.08 *m/z*. Similar trends were noticed for multiple analytes and hypothetical degradation reactions are proposed (FIGURE 6). Additional proposed reactions not shown in Figure 6 include 709.14 *m/z* to 357.06 *m/*z and Spinatoside-4’-β-D-(2′-O-feruloyl-glucuronide) (697.14 *m/z*) to Patuletin (331.05 *m/z*). While the relative bioaccessibility estimates reported here are quite large for some aglycones, these analytes are among the least abundant flavonoids in raw spinach and found in nanomolar concentrations. Small increases in these molecules result in relatively large percent increases. While these data suggest a simultaneous over and slight under estimation of some analytes, *in vitro* digestions still represent an invaluable tool to contextualize the potential health benefits of a food or food product. Studies leveraging isolated molecules and potentially intrinsic labeling would be necessary to elucidate their true absorption, distribution, metabolism, and excretion. Regardless, estimates provided here serve as the most comprehensive profiling of spinach flavonoids pre and post *in vitro* digestion to date and will help breeders and nutrition scientists make data-driven decisions about this crop.

**Figure 6.**
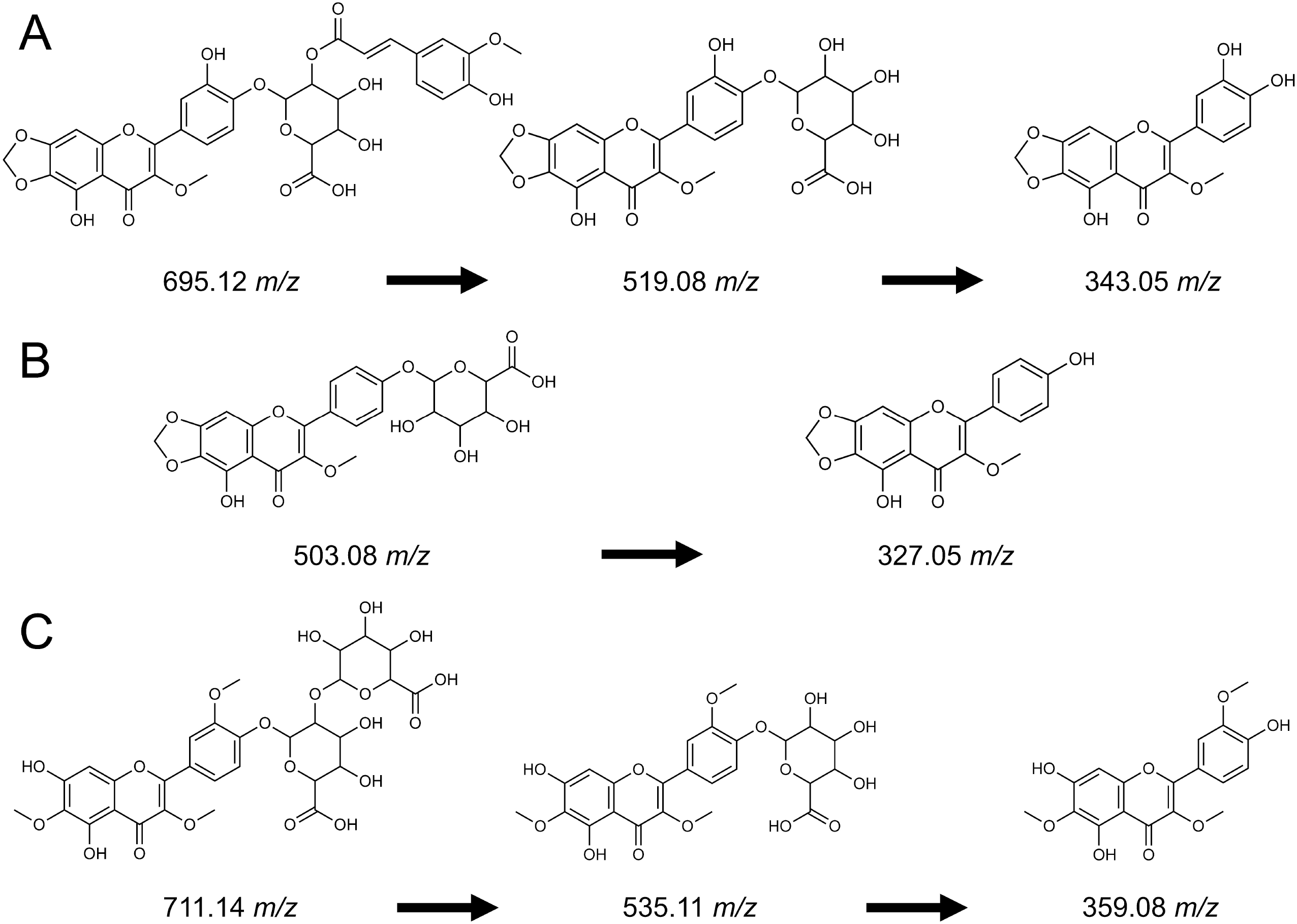
Proposed reactions that occurred during the *in vitro* digestion justifying >100% relative bioaccessibility for multiple spinach flavonoid aglycones not normally seen in high abundance in raw material.

## 5. Conclusions

Here, we reported the range and concentrations of flavonoids in a diverse population of spinach, estimated relative and absolute bioaccessibility of these molecules, and calculated broad sense heritability for these traits. These data were generated using a novel mass spectrometry methodology that allows for comprehensive view of the spinach flavonoid pathway. These data address critical gaps in knowledge about the profiles and bioaccessibility of these molecules. Our data suggest that there may be at least one split in the spinach flavonoid biosynthetic pathway whereby several methylenedioxyflavone species compete for common precursors required for the biosynthesis of other molecules.

We found that absolute bioaccessibility is more heritable than raw material content in the context of spinach flavonoids and will be of interest to plant breeders seeking to improve the phytochemical content of spinach. Indications that environmental factors play a major role in both flavonoid biosynthesis and bioaccessibility provide numerous opportunities for the CEA community to develop environmental manipulation strategies that could alter the phytochemical profile and potential health benefits of spinach. Additionally, we discovered that several spinach flavonoids appear to be susceptible to chemical transformation during digestion, which would be enhanced *in vivo* due to brush boarder enzymes and microbiome activity. Many opportunities exist to utilize genomic and metabolomic tools to fully characterize this pathway, define genetic and environmental cues that regulate this pathway, and define how these molecules impact human health.

## Supplementary Materials

The following supporting information can be downloaded at: https://www.mdpi.com/article/doi/s1

## Author Contributions

Conceptualization, M.D.; methodology, M.D., A.T., E.L., B.R., and G.D.; software, M.D. and R.D.; validation, M.D.; formal analysis, M.D.; investigation, M.D., R.D., A.T., and E.L.; resources, M.D.; data curation, M.D.; writing—original draft preparation, M.D.; writing—review and editing, M.D., A.T., E.L., R.D., B.R, and G.D.; visualization, M.D. and R.D.; supervision, M.D.; project administration, M.D.; funding acquisition, M.D. All authors have read and agreed to the published version of the manuscript.

## Data Availability Statement

The original contributions presented in this study are included in the article/supplementary material. Further inquiries can be directed to the corresponding author.

## Acknowledgments

We greatly appreciate the advice from David Brenner and the USDA-ARS GRIN as well as Dr. Massimo Iorizzo at North Carolina State University for assistance with developing the spinach population used in this study. We thank Brian Chlouber for helping culture and harvest plant material.

## Author Disclaimer

The findings and conclusions in this publication are those of the authors and should not be construed to represent any official USDA or U.S. Government determination or policy. Mention of trade names or commercial products in this publication is solely for the purpose of providing specific information and does not imply recommendation or endorsement by the U.S. Department of Agriculture. The USDA is an equal opportunity provider and employer.

## Abbreviations

The following abbreviations are used in this manuscript:

BLUE: Best linear unbiased estimate
CEA: Controlled environment agriculture
FW: Fresh weight
LED: Light emitting diode
UHPLC-MS/MS: Ultra-high performance liquid chromatography mass spectrometry
REML: Restricted maximum likelihood
SPE: Solid phase extraction

